# Dachsous is a key player in epithelial wound closure by modulating cell shape changes and cytoskeleton dynamics

**DOI:** 10.1101/2024.07.17.603948

**Authors:** Patrícia Porfírio-Rodrigues, Telmo Pereira, Antonio Jacinto, Lara Carvalho

**Author notes:** Instituto Gulbenkian de Ciência, Oeiras, Portugal (Patrícia Porfírio-Rodrigues).

## Abstract

Epithelia are vital tissues in multicellular organisms, acting as barriers between external and internal environments. Simple epithelia, such as those in embryos and the adult gut, have the remarkable ability to repair wounds efficiently, making them ideal for studying epithelial repair mechanisms. In these tissues, wound closure involves the coordinated action of a contractile actomyosin cable at the wound edge and collective cell movements around the wound. However, the dynamics of cell-cell interactions during this process remain poorly understood.

Here, we demonstrate that Dachsous (Ds), an atypical cadherin associated with Planar Cell Polarity, is crucial for efficient epithelial repair in the *Drosophila* embryonic epidermis. We show that the absence of Ds leads to delayed wound closure, impaired actomyosin cable formation, and altered cell shape changes. Additionally, we reveal that Occluding Junctions are necessary for the proper apical localization of Ds, suggesting an unanticipated interaction between these two molecular complexes. This study identifies Ds as a novel player in epithelial repair and highlights the need for further investigating the molecular mechanisms by which Ds modulates cell shape and tissue morphogenesis.

**Summary statement:** This study shows that the atypical cadherin Dachsous is essential for epithelial wound closure, influencing cytoskeletal dynamics and cell morphology, with Occluding Junctions regulating its subcellular localization.

## Introduction

Simple, one-layered epithelia can repair their wounds in an efficient and scarless manner, being thus bona-fide models to study epithelial repair mechanisms (Eming et al., 2014; Lim et al., 2024; Zulueta-Coarasa and Fernandez-Gonzalez, 2017). In these tissues, wound closure is promoted by a contractile, supracellular structure assembled at the wound edge formed by F-actin and non-muscle Myosin 2 (Myo2) – the so-called actomyosin cable – that quickly pulls cells together to close the gap (Bement et al., 1993; Martin and Lewis, 1992; Wood et al., 2002; Zulueta-Coarasa and Fernandez-Gonzalez, 2017). Simultaneously, cells rearrange and move collectively without losing cell-cell contacts and tissue integrity (Razzell et al., 2014; Tetley et al., 2019). In *Drosophila* embryonic wounds, remodelling of Adherens (AJs) and Occluding Junctions (OJs) at the wound edge is required for actomyosin cable assembly and efficient closure (Abreu-Blanco et al., 2012; Carvalho et al., 2014; Carvalho et al., 2018; Hunter et al., 2015; Matsubayashi et al., 2015; Wood et al., 2002). However, whether these junctions play a relevant role in the cellular rearrangements and shape changes during the repair process remains poorly understood.

Cell and tissue polarity play essential roles in epithelial morphogenesis and disease. In epithelial tissues, cells are polarized along their apicobasal axis, displaying functionally and molecularly distinct apical, lateral, and basal cortical domains (Buckley and St Johnston, 2022; Stephens et al., 2018; Tepass, 2012). At the tissue level, epithelia also present asymmetries in the plane orthogonal to the apicobasal axis, referred to as planar cell polarity (PCP) (Butler and Wallingford, 2017; Devenport, 2014). Dachsous (Ds) and Fat (Ft) are evolutionarily conserved atypical cadherins known for their roles in regulating PCP in diverse§ tissues (Sharma and McNeill, 2013). Ds and Ft are asymmetrically localized within cells and bind to each other in a heterophilic fashion. This molecular asymmetry can lead to a wide range of collective cell and tissue behaviours, such as the positioning of specific cellular structures (e.g. hairs, cilia), the orientation of cell divisions, and the regulation of cell shape changes and collective cell migration in various organisms, from *Drosophila* to mammals (Blair and McNeill, 2018; Fulford and McNeill, 2020). Importantly, disruptions in the Ds-Ft pathway are linked to several developmental defects and diseases, including rare genetic disorders and cancer (Alders et al., 2014; Cappello et al., 2013; Kasiah and McNeill, 2023; Pilehchian Langroudi et al., 2017). However, the molecular and cellular mechanisms by which these conserved cadherins modulate cell shape and tissue morphogenesis during development and disease remain unclear.

Here, we show that Ds is essential for efficient wound closure in the *Drosophila* embryonic epidermis by influencing actomyosin cable formation and cell shape changes and rearrangements. Furthermore, we reveal that OJs modulate the proper apical distribution of Ds protein in these cells. As OJs are known to be required for cell shape changes during wound closure, this work uncovers a novel mechanism by which OJs and Ds might interact during epithelial wound closure.

## Results and Discussion

### Ds is required for efficient wound closure

To address the role of the Ds-Ft polarity complex in wound closure in the *Drosophila* embryonic epidermis, we examined wound closure dynamics in mutants for different Ds-Ft pathway components. We wounded the ventral epidermis of late embryogenesis embryos (stage 15) expressing an F-actin reporter under the control of a ubiquitous promoter (*da*-Gal4>UAS-Cherry::Moesin-ABD) (Millard and Martin, 2008) using a UV laser, and imaged wound closure dynamics by time-lapse confocal microscopy. We found that while wild-type (WT) embryos closed their wounds in 67±16.7 min, *ds-/-* mutants took almost 99±40.4 min (average±SD), representing a 48% increase in the duration of wound closure (Fig. 1A,B,D,E; Movie 1). Conversely, *ft*-/-mutants closed their wounds in a similar average time as WT, albeit with higher variability (70±23.4 min; Fig. 1A,C,D,E; Movie 1). We also examined mutants for two other Ds-Ft pathway components: Four-jointed (Fj) and Dachs. Fj is a Golgi kinase that phosphorylates Ft and Ds, modulating their binding affinity (Ishikawa et al., 2008), while Dachs is an unconventional myosin acting as an effector of the Ds-Ft pathway, binding both Ds and Ft (Blair and McNeill, 2018; Mao et al., 2006). Interestingly, *fj-/-* mutants showed a similar phenotype to *ds-/-* mutants, taking 110±27.0 min to complete wound closure (Figs 1E, S1A,C; Movie 2). In contrast, *dachs*-/-mutants closed their wounds with similar timing as WT, though higher variability (68±37.9 min; Figs 1E, S1B,C; Movie 2). These results suggest that Ds and Fj play important roles in epithelial wound closure, whereas Ft and Dachs appear to be less essential for this process.

**Fig. 1.**
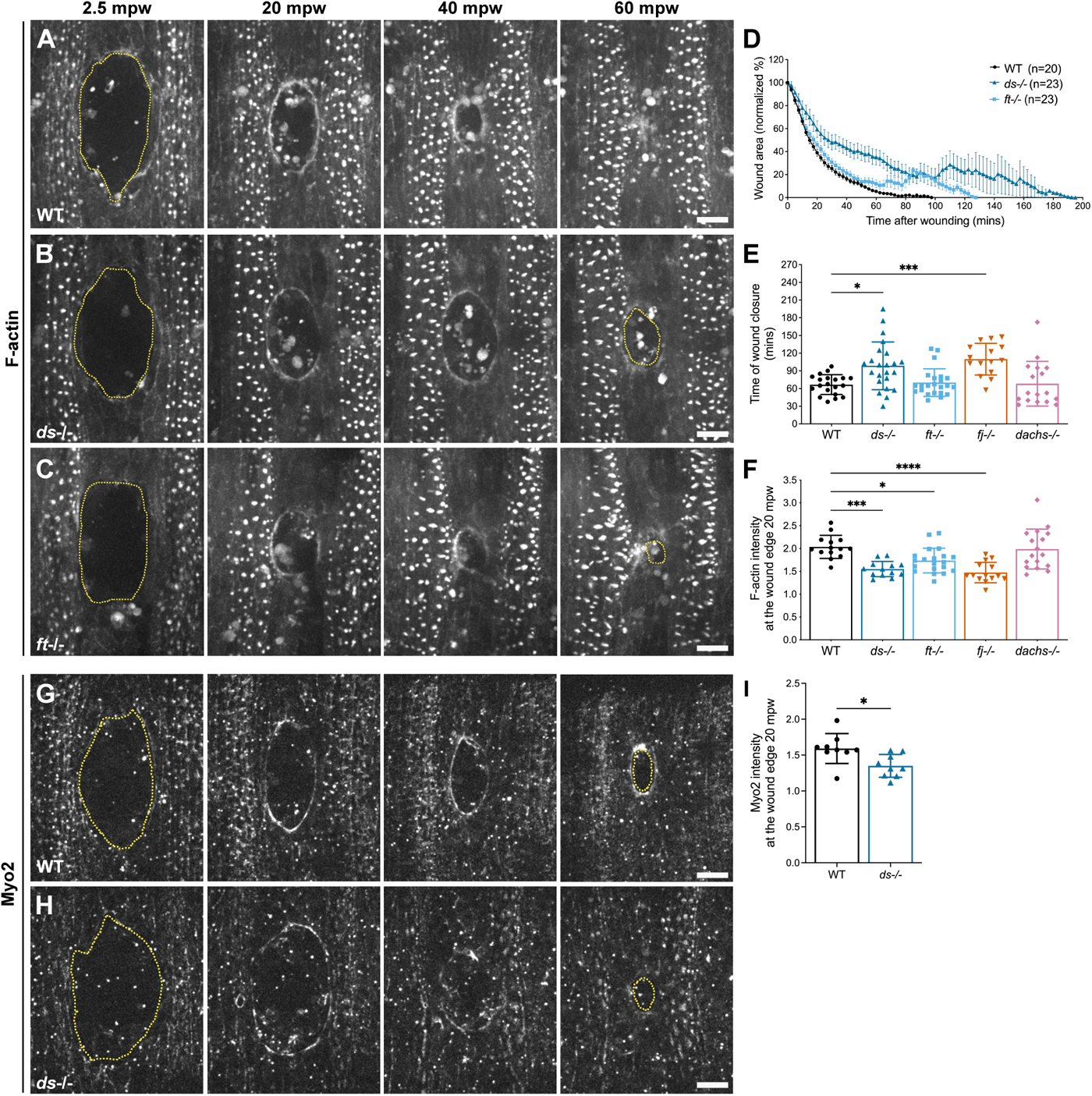
Dachsous is required for wound closure and actomyosin cable formation. (A-C) Embryos expressing mCherry::MoesinABD depicting F-actin at 2.5, 20, 40 and 60 min post wounding (mpw) in WT (A), *ds-/-* (B) and *ft*-/- (C) mutants. (D-E) Graphs showing wound area over time in WT, *ds-/-* and *ft-/-* mutants (D) and time of wound closure in WT (n=20), *ds-/-* (n=23), *ft-/-* (n=23), *fj-/-* (n=15) and *dachs-/-* (n=17) mutants (E). (F) Graph showing F-actin intensity at the wound edge at 20 mpw in WT (n=14), *ds-/-* (n=13), *ft-/-* (n=20), *fj-/-* (n=14) and *dachs-/-* (n=17) mutants. (G,H) Embryos expressing sqh-sqh::mKate depicting Myo2 at 2.5, 20, 40 and 60 mpw in WT (G) and *ds-/-* mutants (H). (I) Graph showing Myo2 intensity at the wound edge at 20 mpw in WT (n=9) and *ds-/-* mutants (n=9). Images in A-C,G,H are maximum z projections; anterior is to the left; yellow dashed line marks the wound edge; scale bars:10 µm.

To gain further insights into the mechanisms through which the Ds pathway functions in wound closure, we analysed the formation of the actomyosin cable by measuring F-actin levels at the wound edge. We observed that both *ds*-/- and *fj-/-* mutants showed a significant reduction in F-actin levels at the wound edge at 20 min post-wounding compared to WT (24% reduction for *ds-/-*, 29% reduction for *fj-/-;* Fig. 1F). Interestingly, *ft-/-* mutants also exhibited a slight decrease in F-actin levels (17%) whereas *dachs-/-* mutants showed levels similar to controls (Fig. 1F). These F-actin phenotypes seem to be specific to the actomyosin cable and wound closure, as F-actin levels in epidermal cells before wounding were similar among all genotypes (Fig. S1D). Supporting the hypothesis that Ds influences actomyosin cable formation, Myo2 levels at the wound edge were also decreased in *ds-/-* mutants compared to WT (Fig. 1G,H,I, Movie 3). Additionally, as for F-actin, Myo2 intensity in cells before wounding was similar between both genotypes (Fig. S1E), indicating that Ds is specifically required for Myo2 dynamics at the wound edge.

Altogether, these results suggest that the Ds-Ft complex contributes to proper wound closure and actomyosin cable formation, with Ds playing a more relevant role than its partner Ft. Although other transmembrane binding partners for Ds have not been identified, previous studies suggest that Ds might may bind to ligands other than Ft (Matakatsu and Blair, 2006). Several studies also report that the loss-of-function phenotypes of Ds and Ft can differ as the intracellular domains of these proteins bind to different intracellular molecules, resulting in diverse and sometimes opposite cellular behaviours (Blair and McNeill, 2018). Additionally, our observation that *dachs*-/-mutants show a milder wound closure phenotype than *ds-/-* and *fj-/-* mutants suggests that Ds acts through alternative effector molecules. Future studies should aim to identify further downstream targets of Ds and additional extracellular binding partners.

### Ds regulates cell shape changes during wound closure

During *Drosophila* wound closure, cells change shape, intercalate and stretch towards the wound (Razzell et al., 2014; Tetley et al., 2019). To investigate whether Ds influences these processes, we performed time-lapse imaging in WT and *ds-/-* mutants marked with E-cadherin-GFP (Huang et al., 2009), which highlights cell boundaries (Fig. 2A,B, Movie 3). In the embryonic epidermis, cells positioned dorsoventrally to the wound (DV cells) typically shorten in the DV axis, while cells lying anteroposteriorly (AP cells) tend to stretch along the DV axis (Razzell et al., 2014). Thus, we measured the DV elongation ratios of cells adjacent to the wound and calculated the changes in these ratios at various time points during wound closure compared to pre-wounding. Interestingly, we observed that DV cells became less elongated at the beginning of wound closure and more stretched towards the end, both in WT and *ds-/-* mutants (Fig. 2C). Notably, DV cells in *ds-/-* mutants showed significantly greater elongation ratios at the end of closure compared to WT (Fig. 2C). Conversely, AP cells showed similar behaviours in both WT and *ds-/-* mutants, becoming highly elongated shortly upon wounding and then gradually decreasing their DV elongation ratios during wound closure (Fig. 2D). These findings suggest that, in the absence of Ds, a subset of cells undergoes significant deformation towards the end of closure. Given the extensive cell shape changes observed at the wound edge, we hypothesized that apical cell areas might also be affected. To test this, we measured changes in apical cell areas during wound closure. In both WT and *ds*-/-mutants, cells adjacent to the wound edge rapidly decreased their areas post-wounding compared to their pre-wound areas (Fig. 2F,G). Interestingly, DV cells in *ds*-/-mutants showed a more pronounced decrease in apical area than WT cells, as early as 2.5 min post-wounding, suggesting that the absence of Ds leads to a different cellular response to the wound.

**Fig. 2.**
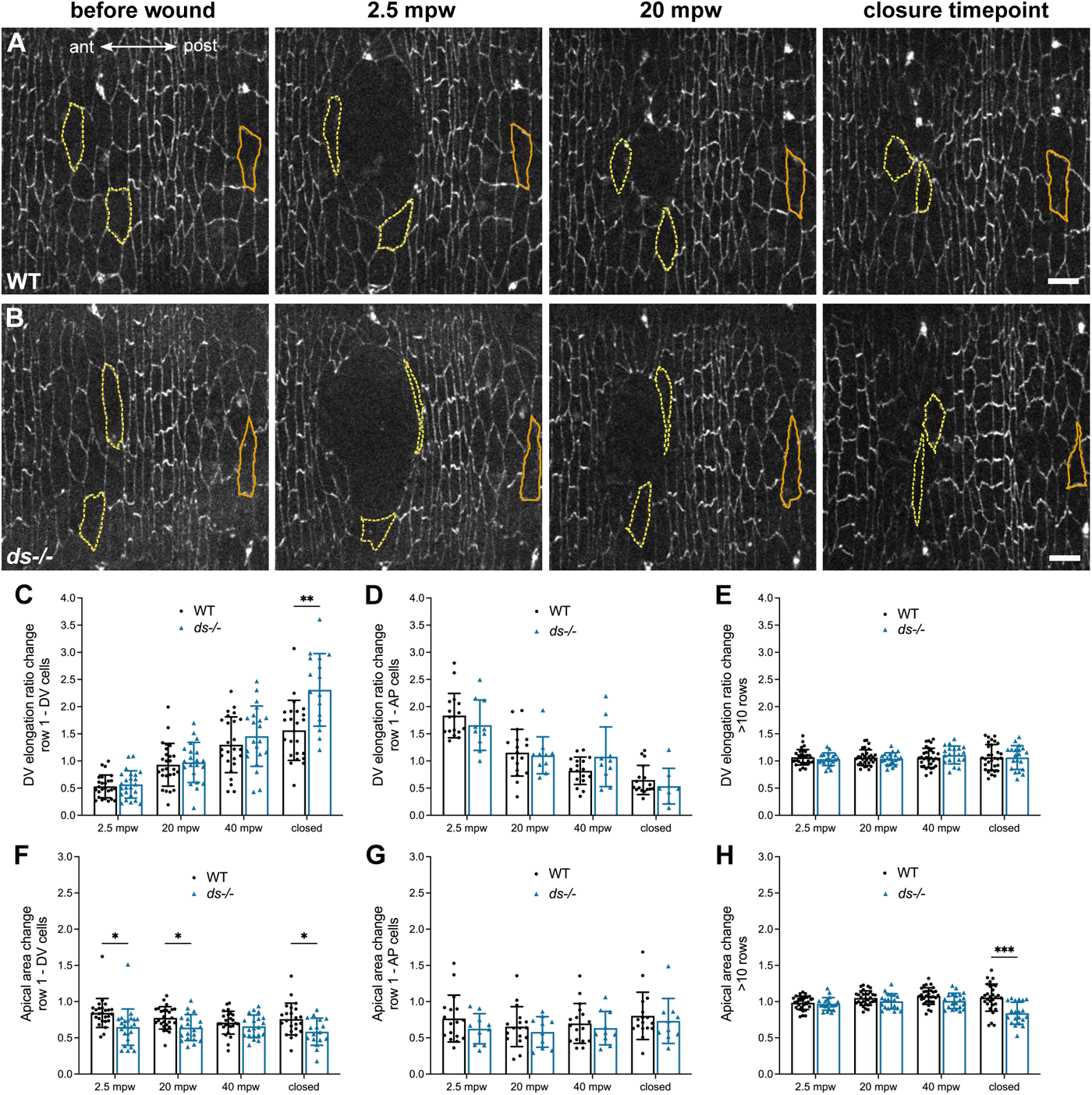
Dachsous influences cell shape changes during wound closure. (A,B) Embryos expressing E-cad::GFP to show cell shapes immediately before wounding, at 2.5 and 20 post-wounding (mpw), and at the time point of wound closure in WT (A) and *ds-/-*mutants (B). Cells outlined with yellow dashed lines are adjacent to the wound edge, cells outlined with orange full lines are 10 rows or further away from the wound. (C-H) Graphs showing DV elongation ratio change (C-E) and apical area change (F-H) in cells lying at the DV (C,F) or AP wound edge (D,G), and in cells at least 10 rows further from the wound (E,H) in WT (n=17-30 cells from 6 embryos) and *ds-/-*mutants (n=10-23 cells from 4 embryos). Images in A,B are maximum z projections; anterior is to the left; scale bars: 10 µm.

To determine if these effects on cell shape changes extend to cells distant from the wound, we examined cells 10 or more cell rows away from the wound edge. While their DV elongation ratios were unchanged during wound closure in both WT and *ds-/-* mutants (Fig. 2E), apical cell areas showed significant alterations (Fig. 2H). Whereas WT cells maintained their apical areas unchanged throughout wound closure, *ds-/-* mutant cells significantly decreased their areas towards the end of closure when compared to WT cells.

Altogether, these results suggest that Ds deficiency alters cell shape changes during wound closure, even in cells further from the wound edge.

Previous studies have shown that cells close to the wound undergo junctional remodelling and exchange neighbours (Carvalho et al., 2018; Razzell et al., 2014; Tetley et al., 2019). To address whether Ds influences this process, we calculated the frequency of neighbour exchange events among cells adjacent to the wound edge in *ds*-/-mutants and WT. Interestingly, *ds-/-* mutants exhibited fewer neighbour exchange events than WT (Fig. S2A-C), indicating that cellular rearrangements are altered in the absence of Ds.

Altogether, these results suggest that Ds influences cell shape changes and rearrangements during wound closure, further supporting our hypothesis that this protein plays an essential role in the wound closure process.

### Ds influences epidermal cell morphology

Considering that Ds appears to influence cell shape changes during wound closure, we wondered whether Ds also regulates cell morphology in the uninjured epidermis. In late embryogenesis, the *Drosophila* epidermis is composed of two cell types: smooth cells and denticle-producing cells. Despite showing distinct cell shapes, both cell types exhibit planar polarity, being elongated along the DV axis and forming aligned columns of cells (Price, 2006; Walters et al., 2006). While Ft is known to regulate planar polarity and junctional remodelling in these cells, particularly in denticle-producing cells (Marcinkevicius and Zallen, 2013), the role of Ds in these processes remain unexplored.

To address this, we imaged the uninjured epidermis of WT and *ds-/-* mutants marked with E-cadherin-GFP and measured both DV elongation and apical cell area of smooth and denticle cells across different columns. As expected, in WT, smooth cells were generally wider in their AP axis and larger than denticle cells, reflected in lower DV elongation ratios and higher apical cell areas (Fig. 3A,B,E,F). When comparing WT and *ds-/-* mutants, we observed that both smooth and denticle cells exhibited similar DV elongation ratios in both genotypes, except for cells in denticle column 6, which showed a significant increase in their DV elongation ratio in *ds-/-* mutants compared to WT (Fig. 3C,D,E). Regarding apical cell areas, smooth cells were significantly larger in *ds-/-* mutants than in WT, particularly in smooth cell columns 2 and 4 (Fig. 3F).

**Fig. 3.**
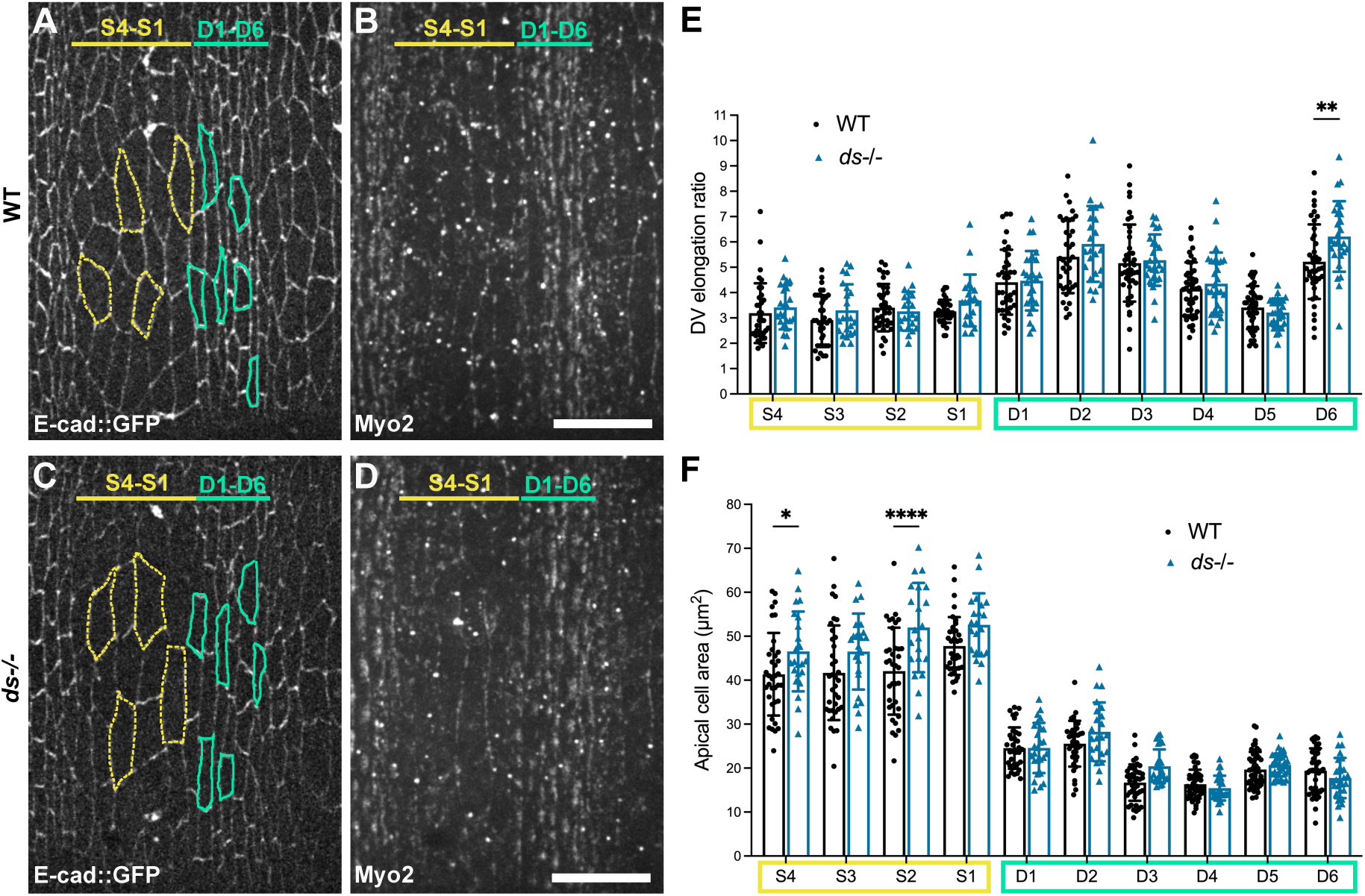
Dachsous influences epidermal cell morphology. (A,C) Morphology of smooth (outlined in yellow dashed line) and denticle cells (outlined in green full line) marked by E-cad::GFP in WT (A) and *ds-/-* mutants (C). (B,D) Myo2 localization allows the identification of smooth (S1-S4) and denticle cells (D1-D6). (E,F) Graphs showing DV elongation ratio (E) and apical cell area (F) in WT (n=36-49 cells from 9 embryos) and *ds-/-* mutants (n=21-32 cells from 6 embryos). Images in A-D are maximum z projections; anterior is to the left; scale bars: 10 µm.

These results suggest that Ds modulates the morphology of embryonic epidermal cells, potentially influencing their response to wounding.

### Occluding Junctions regulate Ds apical-basal distribution

The cell morphology phenotypes observed in *ds-/-* mutants resemble the OJ loss-of-function phenotypes previously described in both uninjured and injured *Drosophila* embryonic epidermis (Carvalho et al., 2018). Specifically, OJ and *ds-/-* mutants show a similar increase in apical area of smooth cells before wounding and impaired cellular rearrangements during wound closure. We thus hypothesized that OJs might be regulating Ds function. To test this we investigated whether the absence of OJs influences Ds expression and localization in epidermal cells. Using a validated Ds::GFP knock-in transgenic line (Brittle et al., 2012), we examined the distribution of endogenous Ds in the epidermis of WT and mutants for the OJ core component Kune (Nelson et al., 2010; Oshima and Fehon, 2011). As expected (Lawlor et al., 2013), Ds was differentially localized in the denticle field in WT, being highly enriched at the plasma membranes between columns 4, 5 and 6 (Fig. 4A,B). In smooth cells, Ds was enriched in the plasma membranes of most cells, except at the boundary between the first column of smooth cells and the first column of denticle cells (Fig. 4A,B). In *kune-/-* mutants, Ds::GFP was less enriched at the plasma membrane and more localized in the cytoplasm compared to WT (Fig. 4A,C). Indeed, the plasma membrane to cytoplasm ratio of Ds::GFP fluorescence significantly decreased in *kune-/-* mutants compared to WT at the boundaries between smooth cell columns 3 and 4, and between denticle cell columns 4 and 5 (Fig. 4E). This suggests that functional OJs are necessary for proper Ds localization at the plasma membrane of smooth and denticle cells.

**Fig. 4.**
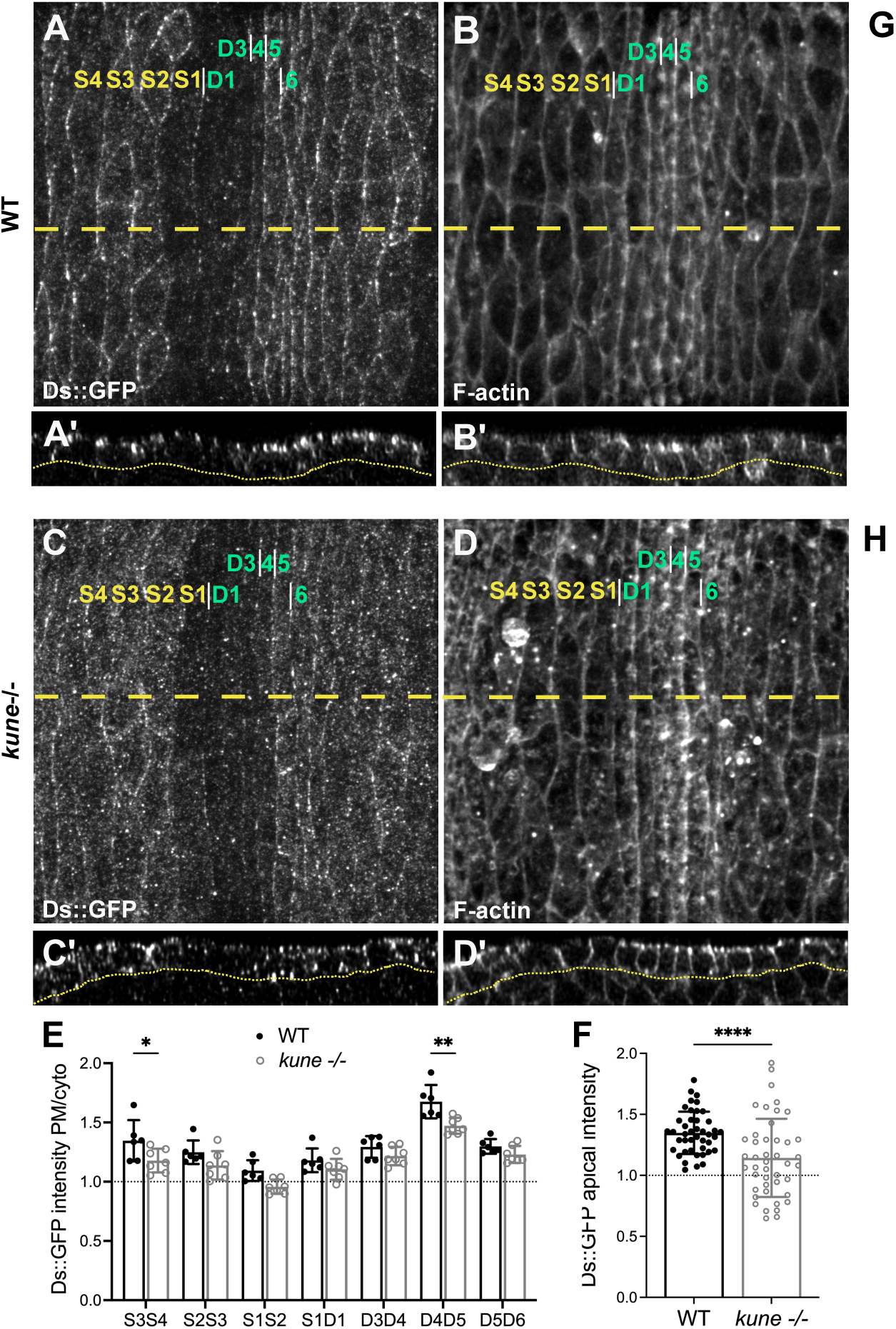
Occluding Junctions are required for proper localization of Dachsous. (A-D) Localization of endogenous Ds::GFP protein (A,C) and F-actin (B,D) in WT (A,B) and *kune-/-* mutants (C,D). S1-S4 mark smooth cell columns, D1-D6 mark denticle cell columns, identified by their distinctive F-actin pattern. Dashed lines in (A-D) mark the localization of longitudinal sections shown in (A’-D’), representing apical-basal views. Dashed lines in (Á-D’) outline the boundary between the basal side of the epidermis and the underlying tissue. Images in A-D are maximum z projections; images in A’-D’ are xz sections (apical is up); anterior is to the left; scale bars:10 µm. (E,F) Graphs showing Ds::GFP intensity at the cell boundaries measured in maximum z projections (n[WT]=6 embryos, n[*kune-/-*]=7 embryos) (E) and Ds::GFP apical intensity ratios measured in longitudinal sections (n[WT]=43 cells from 6 embryos, n[*kune-/-*]=44 cells from 7 embryos) (F). (G,H) Schematic model of how OJs might influence Ds localization and wound closure.

To further explore this, we investigated the subcellular localization of Ds in these cells. As Ds typically localizes at the apical side of epithelial cells, just above AJs (Ma et al., 2003), we analysed whether the apical-basal distribution of Ds was affected in *kune-/-* mutants. We observed that, while in WT Ds accumulated at the apical side of most epidermal cells (Fig. 4A’,B’), in *kune-/-* mutants Ds was more evenly distributed along the apical-basal membrane and cytoplasm (Fig. 4C’,D’). To quantify the apical localization of Ds, we measured the Ds::GFP levels at the apical region and normalized them to the total plasma membrane fluorescence of Ds::GFP. We detected a significant decrease in apical Ds::GFP in *kune-/-* mutants compared to WT (WT=1.4 vs *kune*=1.1, Fig. 4F), with almost half of the mutant cells showing an apical ratio of Ds::GFP < 1.0, indicating that Ds can even accumulate at the basal instead of the apical region in the absence of OJs. These results suggest that OJs regulate the proper apical localization of Ds in embryonic epidermal cells. To our knowledge, this is the first report describing a potential regulator of the apical localization or recruitment of Ds.

OJs have been implicated in epithelial wound closure in *Drosophila* and proposed to regulate mechanical properties and cellular rearrangements (Carvalho et al., 2018). Additionally, the loss of function of a single OJ core component is sufficient to induce the disassembly of the entire OJ multiprotein complex, leading to mislocalization of other OJ components and their spreading along the lateral plasma membrane (Izumi and Furuse, 2014; Rice et al., 2021).

Therefore, the loss of OJs may prevent the proper recruitment of Ds to the apical membrane or promote the ectopic localization of Ds throughout the lateral plasma membrane (Fig. 4G,H). Future studies should further address how Ds apical localization is modulated by OJs and other polarity regulators and the relevance of this for Ds function. Interestingly, in *Drosophila* post-embryonic epithelia the loss of OJ proteins can lead to planar polarity phenotypes through yet unidentified mechanisms (Rice et al., 2021), further supporting the hypothesis that OJs might interact with PCP pathways such as Ds.

Altogether, we uncover a novel role for Ds in regulating wound repair in the *Drosophila* embryonic epithelium. We show that Ds is involved in actomyosin cable stabilization and cell shape changes during wound closure, though the precise molecular mechanisms remain unclear. One hypothesis is that Ds-mediated cell adhesion impacts cell contractility. Even before wounding, cells without Ds show increased apical areas, which are typically associated with reduced junctional tension (Tsoumpekos et al., 2018). This impaired cell morphology and potentially altered mechanical properties might underlie the cell shape changes observed during wound closure in *ds-/-* mutants, consequently influencing actomyosin cable formation, as previously proposed for OJs (Carvalho et al 2018). Uncovering new Ds effectors and investigating its interactions with the actomyosin cytoskeleton and OJs will further elucidate the mechanisms by which this pathway contributes to epithelial wound healing and morphogenesis.

## Materials and methods

### *Drosophila* strains and genetics

Crosses were performed at 25°C or 29°C on standard *Drosophila* medium. All strains used were purchased from the Bloomington Drosophila Stock Center (Indiana, USA) unless stated otherwise. Flybase release FB2024_02 was used to find stock and gene information (Jenkins et al., 2022).

Mutant alleles and chromosomal deletions used were: *ds*^05142^, *Df(2L)Exel8003*, *ft*^8^, *ft*^attp2^, *fj*^d1^, *fj*^N7^, *dachs*^GC13^, *Df(2L)Exel7038*, *kune*^C309^. All alleles and deletions were crossed to balancer stocks that express GFP under the control of the *twist* promoter (Halfon et al., 2002) and homozygous mutant embryos were identified by the absence of GFP fluorescence. Transallelic combinations or combinations with chromosomal deletions were used whenever possible to avoid the effects of secondary mutations. The *w*^1118^ strain was used as a control (referred to as WT). For confocal imaging, mutant alleles and deletions were recombined with live reporter lines.

The live reporters used were: *UAS-Cherry::MoesinABD* (Millard and Martin, 2008) under the control of the ubiquitous promoter *da-Gal4* (Wodarz et al., 1995) to mark F-actin; *sqh::3xmKate2* to mark endogenous Non-muscle Myosin 2 light chain (Pinheiro et al., 2017); *E-cad::GFP* (Huang et al., 2009) to mark endogenous E-cadherin; and *Ds::GFP* to mark endogenous Ds (Brittle et al., 2012).

### Live confocal imaging

Live imaging was performed as described (Carvalho et al., 2018). Dechorionated stage 15-16 embryos were mounted on their ventral side on glass-bottom culture dishes (MatTek) with embryo glue (double-sided tape diluted in heptane) and covered with halocarbon oil 27 (Sigma). Images were acquired at 25°C on a Nikon/Andor Revolution XD Spinning Disk Confocal Microscope with an EMCCD Camera (iXon 897) using the iQ software (Andor Technology, Belfast, UK), and a 60X Plan Apo VC PFS N.A. 1.4 oil-immersion objective. Individual z slices with a step size of 0.34 µm were acquired as a single time point or every 30 s or every 2.5 min for 60**–**300 min for time-lapses until the point of wound closure.

### Laser wounding

Wounds were inflicted in the ventral embryonic epidermis using a nitrogen laser-pumped dye laser (wavelength 435 nm, Andor Micropoint Galvo) using the microscope setting described in the section above. Each embryo was wounded only once and in a single spot in a smooth field epidermal region.

### Immunohistochemistry and imaging of fixed samples

Embryos of the desired genotype (WT and *kune -/-* mutants positive for Ds::GFP) were collected at stage 15, dechorionated, transferred to a glass vial containing a mix of 1:1 heptane and 4% paraformaldehyde in Phosphate-Buffered Saline (PBS) and fixed for 40 min at room temperature. The vitelline envelope was removed manually, embryos were washed in PBSTT (PBS + 0.1% Tween-20 + 0.1% Triton X-100), then incubated for 1 h in blocking solution (5% Normal Goat Serum + 0.3% Triton X-100 in PBS), followed by primary antibody and probe incubation overnight at 4°C. The primary antibodies used were rabbit anti-GFP (1:1500, #A11122, ThermoFisher) and mouse anti-Fas3 (1:50, #7G10, obtained from the DSHB, created by NICHD of the NIH and maintained at The University of Iowa, Department of Biology, Iowa City, IA 52242). Following incubation, embryos were washed in PBSTT, incubated in secondary antibodies anti-rabbit Alexa 488 and anti-mouse Alexa 647 (1:250, ThermoFisher), and Alexa 568-conjugated Phalloidin (1:100, Invitrogen) for 2 h in blocking solution, washed in PBSTT, and mounted in 80% glycerol, 2% DABCO (Sigma). Imaging was performed on an LSM 980 Confocal Microscope with a 63X Plan Apochromat N.A. 1.3 oil-immersion objective (Zeiss). Images were acquired using the Zen software (Zeiss) and a step size of 0.23 µm.

### Image analysis and quantifications

All images were processed and analysed using Fiji (Schindelin et al., 2012), except when mentioned otherwise.

#### Wound area

Maximum z projections were obtained from Cherry::MoesinABD stacks, an ellipse was drawn along the wound edge, and the area was obtained using the Measure tool. For each embryo, the area was normalized relative to the initial wound area. For consistency, we only considered wounds with an initial area between 500 μm^2^ and 1700 μm^2^.

#### F-actin and Myo2 fluorescence intensity measurements

To measure F-actin and Myo2 intensities at the wound edge, maximum z projections of Cherry::MoesinABD and sqh::3xmKate stacks were used after background subtraction (rolling ball radius=10). The wound edge was outlined using a 5-pixel-wide (∼1 µm) segmented line and the average intensity was obtained using the Measure tool. To obtain intensity values before wounding, for F-actin images, 10 cells per embryo were measured, while for Myo2 the same cells that formed the wound edge upon wounding were measured, identified with the help of the E-cad::GFP channel that highlights cell boundaries. For F-actin measurements, cells containing actin-rich denticle precursor structures were not included in the measurements as they mask the F-actin present at the wound edge and cell cortex.

#### Cell area and DV elongation ratio

Maximum z projections of E-cad::GFP stacks of WT and *ds-/-* mutants acquiring the whole wound closure process, with a time interval of 2.5 minutes, were used. Cells were outlined and the area and aspect ratio were measured using the Measure tool. To obtain the aspect ratio, cells were fitted to an ellipse and the ratio between the length along the DV axis and the length along the AP axis, referred to as the DV elongation ratio, was calculated. For cell measurements during wound closure, the values for each time point were normalized to the values obtained before wounding, to obtain fold changes. Cells at the wound edge were analysed in two separate groups according to their position along the DV and the AP axes (referred to as DV and AP cells, respectively), as the wound displays an ellipsoid shape which leads to different behaviours in these two groups of cells (Razzell et al., 2014). For consistency, in quantifications of cell morphology during wound closure, wounds with an initial perimeter between 100 and 140 µm were considered. Smooth and denticle cells were identified by their position relative to sensory organ precursors, which are located in the smooth cell layer 2, two layers anterior to denticle layer 1 (Marcinkevicius and Zallen, 2013). The position of cells relative to the wound edge was confirmed using the sqh::3xmKate channel that shows the localization of the actomyosin cable.

#### Neighbour exchange events

Neighbour exchange events in WT and *ds-/-* mutants were quantified as described (Carvalho et al., 2018). Maximum z projections of E-cad::GFP stacks acquired during the first 40 minutes after wound closure, with a time interval of 30 s (in the first 15 min) and of 2.5 min (between 15 and 40 min), were used. For consistency, wounds with an initial perimeter between 100 and 140 µm were considered. All cells at the first row adjacent to the wound edge were quantified. For each cell, the number of events was measured (gain or loss of neighbours). This value was normalized to the total number of cells in the first row to obtain the ratio of neighbour exchange events for each embryo. The position of cells relative to the wound edge was confirmed using the sqh::3xmKate channel that shows the localization of the actomyosin cable.

#### Ds::GFP fluorescence intensity measurements

To measure Ds::GFP intensities in the different smooth and denticle cell layers, maximum z projections of Ds::GFP stacks were used. The F-actin channel (marked by Phalloidin) was used to identify the smooth and denticle cell layers as mentioned above for the cell shape measurements, and the Fas3 channel was used to identify the cell boundaries. The AP boundaries between each cell layer (spanning at least 3 cells) were outlined using a 10-pixel-wide (1 µm) segmented line and the average intensity was obtained using the Measure tool. To calculate the relative enrichment of Ds::GFP at the plasma membrane compared to the cytoplasm, in WT and *kune -/-* mutants, the value obtained for each cell boundary was divided by the value obtained for the cytoplasm of the adjacent (posterior) cells.

To measure Ds::GFP intensities along the apical-basal axis, xy sections were used. Deconvolution of the raw stacks was performed using the Huygens software (SVI) to improve signal-to-noise ratio and resolution. The F-actin and the Fas3 channels were used to confirm the apical-basal limits of each cell and to identify the plasma membrane. The whole apical-to-basal region was outlined using a 10 pixel-wide (1 µm) segmented line and the average intensity was obtained using the Measure tool. The same procedure was applied to the apical half of the cell. The intensity at the apical region was calculated as the ratio between the intensity at the apical region and the total intensity for each cell.

#### Statistical analysis

Statistical analysis was performed using GraphPad Prism and Microsoft Excel. Sample sizes are indicated in the respective figure legends.

The normality of the distributions was assessed before evaluating significant differences among groups. Parametric tests were used for normal distributions, and non-parametric tests were used for non-normal distributions.

To evaluate significant differences between two unpaired groups, we used either a Student’s t-test or a non-parametric Mann-Whitney test. To evaluate significant differences among three or more groups, we used a one-way analysis of variance (ANOVA) followed by the Holm-Sidak test for multiple comparisons or the Kruskal-Wallis test followed by the Dunn test for multiple comparisons. A two-way ANOVA with the Geisser-Greenhouse correction followed by a Sidak multiple comparison test was used to test for significant differences between two groups affected by two factors.

In column plots, columns show the mean and error bars show the standard deviation. In time-course plots, error bars represent the standard error of the mean. ns – not significant, * P < 0.05, ** P < 0.01, *** P < 0.001, **** P<0.0001.

## Acknowledgements

We thank David Strutt, Helen Strutt, Eurico Morais-de-Sá and Yohanns Bellaiche for providing fly stocks, the NMS core facilities for their important contributions, particularly the Microscopy and the Fly Facilities, Ana Tavares, Rita Teodoro and Raquel Lourenço for critically reading the manuscript, and the TRI lab for insightful discussions. Stocks obtained from the Bloomington Drosophila Stock Center (NIH P40OD018537) were used in this study.

## Data availability

All relevant data can be found within the article and its supplementary information.

